# Hormesis without evolution: plastic compensatory responses to herbicide drift in *Oxalis stricta*, a common weed of agriculture

**DOI:** 10.64898/2026.01.24.701511

**Authors:** Anah Soble, Elise C Kanefsky, Olive A Tatara, Regina S Baucom

**Affiliations:** Department of Ecology and Evolutionary Biology, University of Michigan, Ann Arbor MI 48109

**Keywords:** Hormesis, herbicide drift, dicamba, tolerance, overcompensation, reproductive allocation, eco-evolutionary dynamics

## Abstract

- Herbicide drift exposes non-target plants to sublethal doses of agrochemicals, yet its ecological and evolutionary consequences remain poorly understood. Although hormesis—defined as stimulatory responses to low doses of otherwise toxic compounds—has been documented following herbicide exposure, it has rarely been evaluated within an evolutionary ecology framework. Here, we integrate concepts of tolerance and overcompensation to examine herbicide-induced hormesis in the common agricultural weed *Oxalis stricta* across two field experiments.
- We exposed replicated maternal lines to a drift-relevant dose of dicamba and quantified growth, reproductive traits, floral allocation, and pollinator visitation. Dicamba drift consistently increased flower production in both years, revealing a robust plastic shift toward reproductive allocation. However, the fitness consequences of increased flowering differed between years: in 2022, increased flowering was associated with higher seed production through indirect, trait-mediated pathways, whereas in 2023 increased flowering did not translate into detectable differences in reproductive output. Structural equation modelling indicated that dicamba effects on reproduction were largely indirect, mediated through correlated trait responses rather than direct stimulation of fitness.
- Despite consistent plastic responses, we detected little genetic variation in the magnitude of hormesis, suggesting limited potential for adaptive evolution of overcompensation. Dicamba drift also reduced individual flower size, indicating a shift toward larger floral displays composed of smaller flowers. These allocation shifts altered pollinator visitation patterns, primarily through changes in flower number, linking herbicide exposure to trait-mediated changes in plant–pollinator interactions.
- Together, our results demonstrate that low-dose herbicide exposure can generate repeatable compensatory responses that reshape ecological interactions, even when evolutionary responses are constrained.

## Introduction

Hormesis, defined as the stimulatory effect of a subtoxic dose of a chemical that is toxic at higher doses (Calabrese *et al*., 2007), has been documented across a wide range of plant stressors, including pollutants and agrochemicals. In plants, hormetic responses are typically expressed as increased growth, reproduction, or altered allocation to specific traits following sublethal exposure to a chemical (Duke *et al*., 2005; Belz *et al*., 2011; Belz & Duke, 2017). Because these responses directly affect traits linked to fitness and species interactions, hormesis has the potential to influence both population dynamics and evolutionary trajectories. However, hormesis has rarely been examined through an evolutionary ecology lens and remains poorly integrated into frameworks linking sublethal stress exposure to trait evolution and eco-evolutionary feedbacks.

Hormesis is likely to be particularly important in agricultural landscapes, where non-target plants are frequently exposed to herbicide drift *via* the off-target movement of agrochemical droplets or vapor following application. Sublethal exposures to herbicides such as glyphosate and ALS inhibitors have been shown to elicit hormetic responses in various weed species (reviewed in Belz *et al*., 2022; Duke *et al*., 2025), with typical responses including an increase in biomass, plant size, or fitness related traits by nine to 145% that of non-exposed controls (Duke *et al*., 2025). Studies of low-dose herbicide exposure also report highly variable effects that depend on species identity, herbicide formulation, and trait measured. Importantly, sublethal exposure can alter reproductive and phenological traits even when overall biomass or reproductive output is unaffected. Because sublethal herbicide exposure can enhance fitness-related traits and alter phenology, hormesis has been hypothesized to contribute to shifts in weed community composition (Belz & Duke, 2014, 2017; Duke *et al*., 2025), the evolution of herbicide resistance (Belz & Duke, 2014; Belz *et al*., 2022), and changes in interactions with pollinators and other community members (Guedes *et al*., 2022; Iriart *et al*., 2022). However, empirical tests linking hormetic trait responses to longer-term evolutionary outcomes remain limited, as does our understanding of how hormetic responses might alter interactions with other community members, such as pollinators.

As such, key questions remain regarding the ecological and evolutionary significance of hormesis. It is unclear whether hormetic responses are consistent across years, whether they vary genetically within species, or whether exposure to low-dose herbicide stress could drive evolutionary change through selection on hormetic responses or underlying allocation strategies. In addition, because hormesis can simultaneously alter multiple traits that contribute to fitness and species interactions–such as growth, reproduction, and floral display–it remains unknown whether sublethal herbicide exposure reshapes ecological outcomes indirectly by modifying how plants interact with pollinators.

These questions can be addressed using a well-developed framework from evolutionary ecology focused on tolerance and overcompensation (Paige, 1987; Ramula *et al*., 2019), which provides a conceptually aligned approach. In this framework, tolerance is defined as the ability to maintain fitness under stress relative to benign conditions, regardless of damage incurred (following Strauss & Agrawal, 1999; Agrawal, 2000; Baucom & Mauricio, 2004; Iriart *et al*., 2022). As illustrated in Fig. 1, tolerance encompasses a range of fitness responses to stress. Undercompensation (Fig. 1A) occurs when fitness declines under stress relative to control conditions, whereas complete tolerance (Fig. 1B) reflects equivalent fitness across environments. Overcompensation (Fig. 1C), in which fitness increases under stress, represents a direct analogue to hormesis, capturing cases where low-dose exposure stimulates growth or reproduction rather than suppressing it. Importantly, genetic lines may also differ in both baseline fitness and the direction or magnitude of their response to stress (Fig. 1D), revealing genetic variation in compensatory strategies and the potential for evolutionary change.

**Figure 1.**
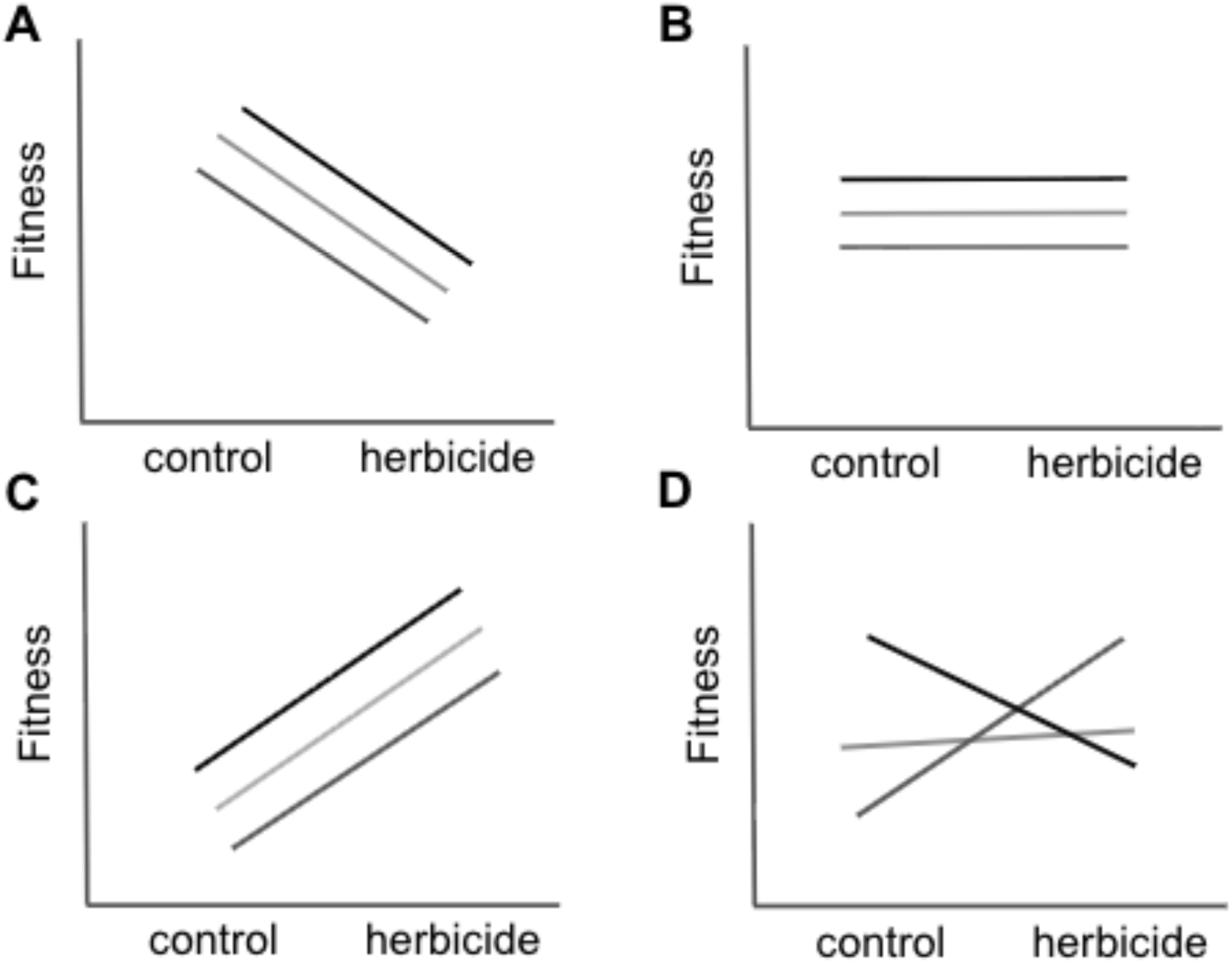
Conceptual illustration of plant fitness responses to sublethal herbicide exposure. Each panel depicts fitness (y-axis) across control and herbicide environments (x-axis) for multiple genetic lines representing maternal or family lines. (A) Undercompensation: all lines show reduced fitness under herbicide relative to control, and lines differ in baseline fitness but not in their response to herbicide. (B) Tolerance: all lines maintain similar fitness across environments, indicating no change in fitness with herbicide exposure. (C) Overcompensation: all lines exhibit increased fitness when exposed to herbicide, again varying in baseline fitness but not in their response. (D) Mixed response: genetic lines differ both in baseline fitness and in the direction and magnitude of their response to herbicide—one line is tolerant, one undercompensates, and one overcompensates—illustrating genetic variation for response to sublethal doses of herbicide.

The extensive literature on overcompensation – most often studied in the context of herbivory damage – has generated a robust foundation linking tolerance, allocation, and evolutionary potential (Paige, 1987; Agrawal, 2000; Ramula *et al*., 2019). Viewed through this lens, hormesis represents a special case of overcompensation in response to chemical stress rather than tissue loss from herbivory. Applying this framework allows hormetic responses to be evaluated as traits with measurable fitness consequences, genetic variation, and potential eco-evolutionary feedbacks under continued exposure to sublethal herbicide stress.

Here, we use the framework of tolerance and overcompensation from evolutionary ecology to examine potential herbicide-induced hormesis in the weed *Oxalis stricta*. Prior greenhouse and field studies show that *O. stricta* survives and often tolerates the herbicide dicamba at drift doses: plants exposed to dicamba drift showed no detectable reduction in growth or flower production (Iriart *et al*., 2022; Baucom *et al*., 2025), although visible injury still occurred (Baucom *et al*., 2025). Our work also suggests overcompensation at drift rates: we previously detected a significant increase in flowering duration following dicamba drift exposure in greenhouse settings (Iriart *et al*., 2022), and unpublished observations show elevated flower production at drift doses of dicamba in the field (Baucom *et al*., 2025). In contrast, the field dose of dicamba (561 g ae ha⁻¹) causes complete mortality in field settings (Baucom *et al*., 2025), consistent with a biphasic, hormetic response. The relevance of variation in hormetic responses has grown with the rapid expansion of dicamba-resistant crop systems in the United States (US EPA, 2021), which has increased off-target drift risk (Soltani *et al*., 2020). As a synthetic auxin herbicide, dicamba disrupts core developmental pathways, and even sublethal exposure may generate context-dependent phenotypes in non-target plants.

We address four related questions. First, does *Oxalis stricta* exhibit increased growth, flower production, or reproductive output under dicamba drift, consistent with overcompensation or hormesis, and are such responses consistent across years? Second, do these responses vary among maternal lines, indicating genetic variation and potential for evolutionary change? Third, does dicamba drift alter patterns of floral allocation, such as trade-offs between flower number and per-flower investment (*e.g.* flower size)? Finally, because herbicide drift may simultaneously influence multiple components of floral display, and because the consequences for pollinator behavior remain poorly understood, we ask whether dicamba drift alters pollinator visitation, and whether any changes are mediated primarily through flower number, flower size, or their combined effects. To integrate hormesis within an evolutionary ecology framework, we test for genotype × environment interactions in fitness and use structural equation modeling to evaluate whether dicamba influences fitness and pollinator visitation directly or indirectly through correlated growth and floral allocation pathways. Together, this approach allows us to assess both the evolutionary potential of hormetic responses and the potential mechanistic links connecting drift exposure to plant fitness and plant–pollinator interactions.

## Materials and Methods

### Study System

*Oxalis stricta* (wood sorrel; Fig. S1) is a wide-ranging, short-lived perennial plant that is native to both North America and Eurasia and is commonly found in agricultural fields, roadsides, cosmopolitan areas, and as herbaceous understory vegetation in temperate forests (Rokhlova, 2013; Imamura & Hariu, 2020). Although *O. stricta* exhibits the morphological remains of a tristylous breeding system, the species is a facultative self-pollinator and is also able to reproduce vegetatively through rhizome production (Doust *et al*., 1985; Marshall, 1987). The species is commonly visited by syrphid flies and honeybees largely for nectar and pollen rewards (*see Results*). Though *O. stricta* mostly self-pollinates it is also able to produce seed when outcrossing and does not undergo apomixis (Doust *et al*., 1981, 1985). The leaves of *O. stricta* contain oxalic acid and are often added to food by humans to impart a sharp flavor (Doust *et al*., 1981, 1985). The species is considered a problematic weed in nurseries, turf, and agriculture (Doust *et al*., 1981, 1985; Marshall, 1987)) and is relatively difficult to remove *via* both mechanical and chemical means (Marble *et al*., 2013).

### Seed Collection

We collected seeds from 10 populations of *Oxalis stricta* growing near soybean or fallow fields in Kentucky and Tennessee, USA (Fig. S1) in the summers of 2018 and 2019 for field experiments. At each collection site, seeds from individual plants were sampled every 1 m along a 20 m transect.

### Field Experiment 1

We performed two field experiments to examine the potential for trait level (*e.g.* increased growth or flowering) responses and how that may or may not lead to overcompensation at the fitness level (*e.g.* seed or fruit production). Throughout this study, we distinguish hormetic responses expressed at the trait level (*e.g.* increased growth or flowering) from overcompensation at the fitness level, recognizing that enhanced trait expression does not necessarily translate into increased reproductive output.

For the first field experiment, on 14 June 2022, we planted replicate seeds from 31 maternal plants sampled from natural populations in a common garden at Matthaei Botanical Gardens (Table S1). Four replicate seeds per maternal line were planted across two blocks, with two treatments (control, dicamba drift) per block (4 replicates × 2 blocks × 2 treatments × 31 maternal lines = 496 individuals). We additionally germinated seeds from each maternal line in a growth chamber in the Biological Science Building’s plant growth facility under an 18 h light : 6 h dark photoperiod at 26°C and replaced non-germinants in the field with transplants on 18 July 2022. Individuals that never germinated or did not survive transplant were excluded from analyses.

After 22 days of growth in the field (9 August 2022), plants were sprayed with either a control or herbicide treatment. Control plants were sprayed with Preference surfactant (non-ionic surfactant blend, WinField Solutions, St. Paul, MN) at 0.1% (v/v). Treatment plants were sprayed with a dicamba solution corresponding to approximately 0.7% of the recommended field dose (561 g ae ha⁻¹), defined based on the equivalent proportion of the labeled field-rate tank concentration and representing a particle drift exposure (Felsot *et al*., 2011), plus 0.1% (v/v) Preference. Plants were sprayed in a single pass until just wet using a handheld multi-purpose sprayer set to a medium-fine mist at an operating pressure of 40–45 PSI. We used a handheld sprayer rather than a CO₂-pressurized sprayer with a constant application rate to better simulate the heterogeneity characteristic of off-target herbicide drift under field conditions. Because applications were made using a handheld sprayer rather than a calibrated boom system, exposure is reported as a proportion of the recommended field dose rather than as an absolute area-based application rate.

We recorded the number of true leaves for each plant immediately prior to herbicide treatment and three weeks post-treatment. We then calculated plant growth as prespray leaf number subtracted from leaf number three weeks post-spray, divided by the number of days between leaf number collection (21). We likewise recorded the number of leaves exhibiting dicamba damage–*i.e.*, leaf cupping or yellowing (Foster & Griffin, 2018)–for an estimate of herbicide damage. We recorded the date of first flowering for each plant (in Julian dates) and recorded the number of open flowers per plant twice a week until the first hard frost, which occurred 17 November 2022. We collected ripe fruits from plants in twice weekly collection rounds beginning in September and continuing until the first frost. We weighed the total seed mass for each individual and the mass of a subsample of 100 seeds from that same individual. We then used this known seed mass to convert total seed weight into an estimated seed count for each individual.

### Field Experiment 2

On 21 April 2023, we planted seeds from 16 maternal lines drawn from eight populations in the greenhouse at Matthaei Botanical Gardens and maintained plants under ambient greenhouse conditions. These maternal lines were randomly selected as a subset of those used in Field Experiment 1 (Table S1). Replicate seeds were obtained either from plants grown in a controlled growth-room environment or from plants grown in the 2022 field experiment, resulting in multiple parental environments represented among experimental individuals. Preliminary analyses indicated that parental environment did not influence overall treatment effects for any trait considered and as such we dropped this effect from the final reported models. Once plants produced at least five true leaves, they were transplanted to the field on 13 June 2023. As in the previous field experiment, we established two environmental blocks, each containing both dicamba and control treatments, and planted between six to twenty-four replicate individuals of each maternal line in every block–treatment combination (16 maternal lines × 2 blocks × 2 treatments × 6-24 replicates (average ∼20 replicates per line) = 689 total individuals).

Approximately three weeks after transplanting (7 July 2023), plants were exposed to the same dicamba and control treatments used previously. We recorded the same vegetative growth measurements as in the prior year and monitored flower production throughout the experiment. Flower number was recorded repeatedly through early September, and flower width, another component of floral display, was recorded one month after spraying (9 August–8 September 2023) by measuring the diameter of five flowers per plant with digital calipers, defined as the distance from the tip of one petal to the tip of the opposing petal. In total, 3,430 flowers were measured; mean flower width per individual was used in subsequent analyses. At the end of the season, we counted fruits per plant and used fruit number as our estimate of fitness, as fruit number was positively correlated with total seed production in the earlier field experiment (r = 0.53, P < 0.001).

Pollinator observations began on 6 September 2023 and were conducted using a subset of experimental plants. Individuals within one block were grouped into spatially proximate clusters of two to ten plants (arrays) that could be observed simultaneously. Ten arrays per treatment were selected, yielding twenty arrays total. Randomly chosen arrays were observed across seven days (rounds) conducted between September and October 2023. General weather patterns were recorded (sunny, partly sunny, cloudy, partly cloudy). Each array was observed for 20 minutes per round; the number of open flowers per plant was recorded at the start of each observation, followed by recording the number of pollinator foraging events, defined as instances in which an insect visitor entered a flower. Floral visitors were categorized into functional groups (honeybee, bumblebee, syrphid fly, wasp, and fly). Across rounds, dicamba-exposed and control plants were observed for a total of 380 and 300 minutes, respectively.

## Data Analysis

### Analysis of plant traits

Field Experiment 1–All analyses were conducted in R (vers 4.3.1; Team, 2025) using R Studio (Posit team, 2024). We analyzed post-spray vegetative growth, date of first flower, total flower number, and seed number from the 2022 field experiment. To improve normality and variance structure, post-spray growth and flowering date were rank-normalized using the orderNorm transformation in the BESTNORMALIZE package (Peterson, 2021), and total flower number was square-root transformed. Seed number, which exhibited strong right skew and heteroscedasticity, was analyzed on its original count scale using a generalized linear mixed-effects model with a Tweedie error distribution and log link, implemented in GLMMTMB (Brooks *et al*., 2017). For post-spray growth, flowering date, and total flower number, we fit linear mixed-effects models using LMER in the LME4 package (Bates *et al*., 2011), with treatment specified as a fixed effect and prespray leaf number included as a covariate where relevant.

In all models, random effects were included to account for experimental blocking and genetic structure. Population and maternal line (nested within population) were included as random intercepts, and block was included as a random intercept to account for spatial structure in the field. To test for genetic variation in treatment responses (*i.e*., maternal-line-specific reaction norms), we evaluated an additional random-effects structure allowing treatment to vary among maternal lines, implemented as a random treatment slope within maternal line (nested within population). Baseline models took the general form:

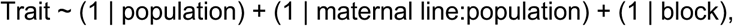

with reaction-norm models additionally including the term (1 + treatment | maternal line:population). Block effects were retained only when they explained non-zero variance. Model fit for all generalized linear mixed-effects models was assessed using the DHARMA package (Hartig, 2024). Fixed-effect significance was assessed with Satterthwaite degrees of freedom using LMERTEST (Kuznetsova *et al*., 2017), and random effects were tested using likelihood ratio tests. Estimated marginal means were determined using EMMEANS (Russell 2025) and back-transformed to the original measurement scales for interpretation.

Field Experiment 2—We applied similar mixed-effects modeling approaches to post-spray vegetative growth, total flower production for the season, flower number during the first month following spraying, and fruit production in the 2023 field experiment. Because many plants had already initiated flowering prior to dicamba drift exposure (*see Results*), we did not analyze treatment effects on the date of first flowering. Models followed the same general structure used for Experiment 1, with treatment specified as a fixed effect and population, maternal line nested within population, and block included as random intercepts.

Continuous response variables (growth and flower number) were analyzed using linear mixed-effects models. Fruit count was analyzed using generalized linear mixed-effects models implemented in GLMMTMB (Brooks *et al*., 2017), with a negative binomial (NB1) error distribution and a log link. Transformations were applied to continuous variables as needed to satisfy assumptions of normality and homoscedasticity. Model fit for all generalized linear mixed-effects models was assessed using the DHARMA package (Hartig, 2024). As in Experiment 1, we evaluated models including a random treatment slope among maternal lines to test for genetic variation in treatment responses.

### Structural equation models of growth- and flower-mediated overcompensation

We used structural equation modeling (SEM) to evaluate whether dicamba exposure influenced reproductive output directly or indirectly through changes in growth and floral allocation. Structural equation modeling allows simultaneous evaluation of direct and indirect effects within an integrated causal framework (Lefcheck, 2016; Grace & Irvine, 2020). We adopted a model-selection approach to SEM (MSA-SEM), comparing alternative causal structures using AIC values from component models fitted relative to a fully saturated model (Garrido *et al*., 2022). As with all structural equation models, these analyses evaluate support for alternative, biologically motivated causal hypotheses rather than establishing definitive mechanistic pathways.

SEMs were fit separately for each field experiment using the PIECEWISESEM package in R (Lefcheck, 2016) to allow for year-specific differences in reproductive traits while maintaining a similar model structure. For both years, dicamba treatment was specified as a predictor variable, with hypothesized effects on post-spray vegetative growth and flower number. Growth was modeled as a predictor of flower production, and reproductive output (seed number in 2022; fruit number in 2023) was modeled as a function of downstream allocation traits. Block was included as a random effect in all component models.

Because reproductive output differed in distribution between years, we used year-specific error structures. In 2022, seed number was modeled using a Tweedie generalized linear mixed model with a log link, whereas in 2023 fruit number was modeled using a negative binomial (NB1) distribution. Growth and flower number were modeled using linear mixed-effects models. Continuous predictors were standardized to facilitate comparison of effect sizes. We manually estimated standardized coefficients for non-Gaussian responses on the link scale, following established recommendations for generalized responses.

For each year, we compared a small set of biologically motivated SEMs that differed in whether reproductive output was mediated by flower number, post-spray growth, or both, and whether direct effects of dicamba on reproduction were included. Model selection was based on tests of directed separation and comparison of AIC values. Full details of candidate models, model comparisons, and goodness-of-fit statistics are provided in the Supplemental Methods.

### Floral allocation given dicamba drift

We analyzed variation in flower width to test whether dicamba drift altered per-flower allocation. Flower width was averaged per individual and square-root transformed to improve normality. Linear mixed-effects models were fit with treatment as a fixed effect, with population and maternal line modeled as random effects. To test whether maternal lines differed in their response to dicamba exposure, we included a random treatment slope among maternal lines, allowing maternal-line-specific reaction norms. The final model structure was:

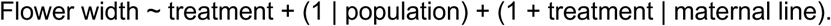

Block was evaluated in preliminary models but excluded because the effect explained negligible variance. Fixed effects were assessed using Satterthwaite-approximated F-tests (LMERTEST; Kuznetsova et al., 2017) and Type III Wald χ² tests (Fox & Weisberg, 2018), and random effects were evaluated using likelihood-ratio tests (Bates et al., 2011). Estimated marginal means were obtained from the final model and back-transformed for presentation.

To test whether dicamba drift altered floral allocation strategies, we modeled average flower width as a function of flower number, treatment, and their interaction using a linear mixed-effects model. Maternal line was included as a random effect. Fixed-effect significance was assessed using Type III tests (Fox & Weisberg, 2018).

### Pollinator visitation

We analyzed pollinator visitation using generalized linear mixed-effects models to account for the non-normal and right-skewed distribution with many zero values. Visitation rate was quantified as the number of pollinator visits per open flower per minute. Models were fit using the GLMMTMB package in R (Brooks *et al*., 2017) with a Tweedie error distribution and a log link function. Dicamba treatment was included as a fixed effect. To account for repeated observations and hierarchical structure in the data, population, observation round, weather category, and experimental array (array × treatment combination) were included as random effects. Model assumptions were assessed using residual diagnostics implemented in the DHARMA package (Hartig, 2024). Fixed effects were evaluated using Type III Wald χ² tests. For interpretation, estimated marginal means were back-transformed to the response scale using the EMMEANS package (Russell 2025), yielding visitation rates expressed as visits per flower per minute with associated 95% confidence intervals.

### Structural equation models of floral trait–mediated pollinator visitation

We used piecewise structural equation modeling (SEM) to evaluate whether dicamba drift influenced pollinator visitation directly or indirectly through changes in floral traits. Models were fit using the PIECEWISESEM package in R (Lefcheck, 2016) and constructed from a set of component models. Dicamba treatment was specified as an exogenous driver with potential direct effects on flower width, flower number, and pollinator visits (visits/min). The goal of this analysis was to determine whether any indirect effects of dicamba on visitation were mediated primarily through changes in individual flower size (flower width) or through changes in floral display size (flower number).

Based on preliminary analyses and tests of directed separation, we evaluated models that differed in whether flower width influenced flower number and whether flower width exerted a direct effect on pollinator visitation. Flower width and flower number were modeled using linear models, while pollinator visit time, which contained many zero values and was strongly right-skewed, was modeled using GLMMTMB (Brooks *et al*., 2017) with a Tweedie error distribution and a log link function. Models were compared using tests of directed separation, Fisher’s C, and summed AIC values to assess support for alternative pathways. For non-Gaussian components, predictor-standardized effects were used to facilitate comparison of effect sizes.

## Results

### Hormetic responses

Dicamba drift induced hormetic responses in some but not all traits (Fig. 2; Table S2). In Field Experiment 1 (2022), growth increased by ∼44% in drift-treated plants (F_1, 171.13_ = 4.62, P = 0.03; Fig. 2A), such that drift-treated plants produced, on average, 4.46 post-spray leaves compared with 3.10 leaves in control plants. Similar to growth, total flower number increased under dicamba drift, rising by approximately 46% (139 vs. 95 flowers in drift vs. control; F_1, 121.01_ = 12.23, P = 0.001; Fig. 2C). The increase in flower production was evident across the majority of sampling dates (Fig. 3A). Flowering phenology (day of first flower), however, showed no difference between treatments (F_1, 106_ = 0.50, P = 0.48; Fig. 2B). Interestingly, we found an ∼18% increase in seed number in treated plants (1.9 x 10^5^ vs. 1.6 x 10^5^ seeds produced on average in drift vs. control; Fig. 2D), but this effect was not significant (F_1_ = 0.58, P = 0.46), indicating that the variance in seed number among individuals was likely high. In fact, the coefficient of variation for seed number in the drift environment was 1.05, whereas the coefficient of variation for flower number was lower (0.58).

**Figure 2.**
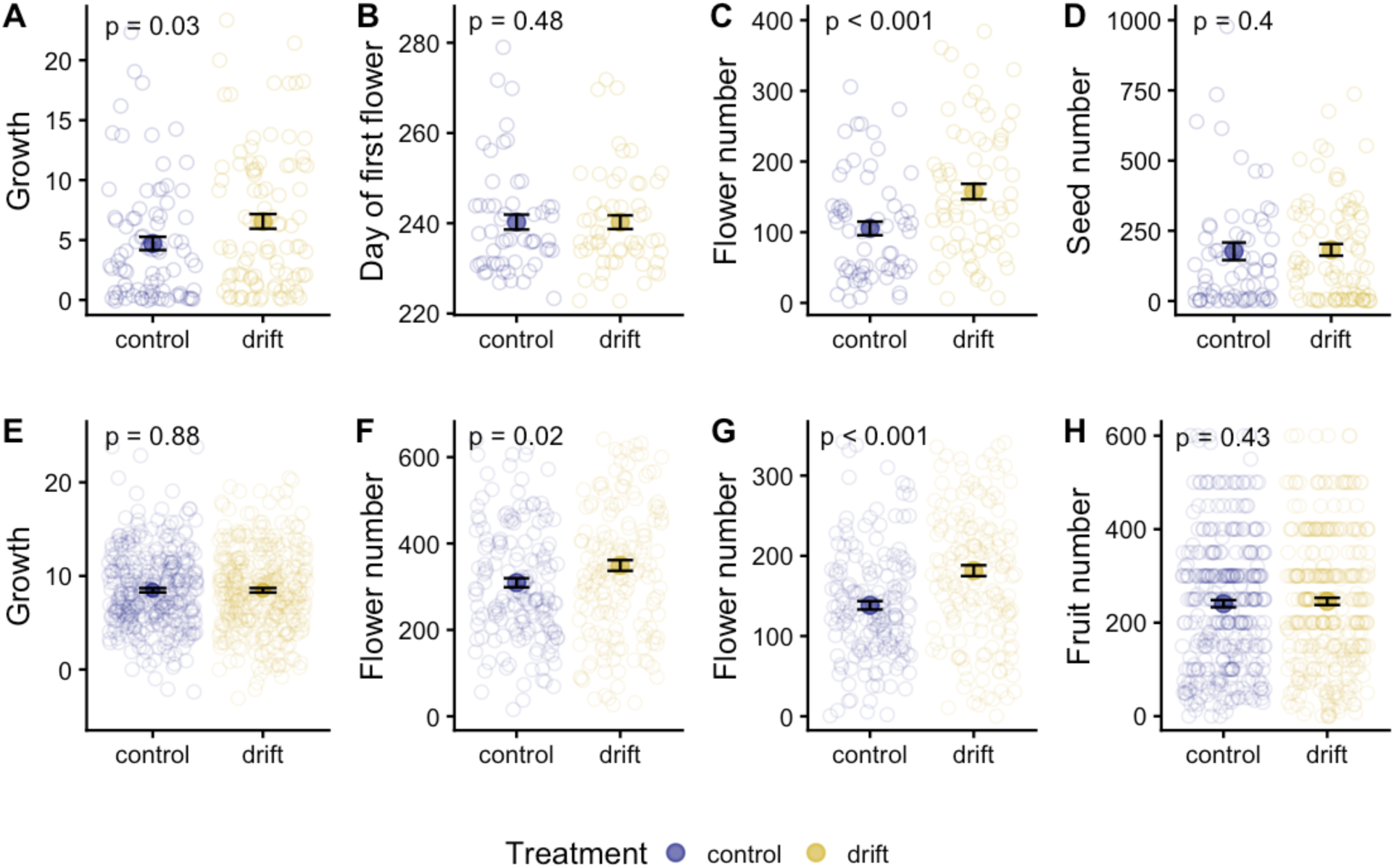
Effects of dicamba drift on plant growth and reproductive traits across two field experiments. Trait means (large, filled circles) ± standard errors (error bars) are shown for control (blue) and dicamba drift–exposed plants (yellow). Individual data points (open circles) represent all measured plants from each experiment. Statistical results (ANCOVA) are provided in Table S1. Panels show: (A) growth rate in Experiment 1 (2022); (B) date of first flowering (Julian day; Exp. 1); (C) total seasonal flower number (Exp. 1); (D) total seed production (thousands; Exp. 1); (E) growth rate in Experiment 2 (2023); (F) total seasonal flower number (Exp. 2); (G) flower number within the first month following dicamba drift exposure (Exp. 2); (H) total fruit number (Exp. 2).

**Figure 3.**
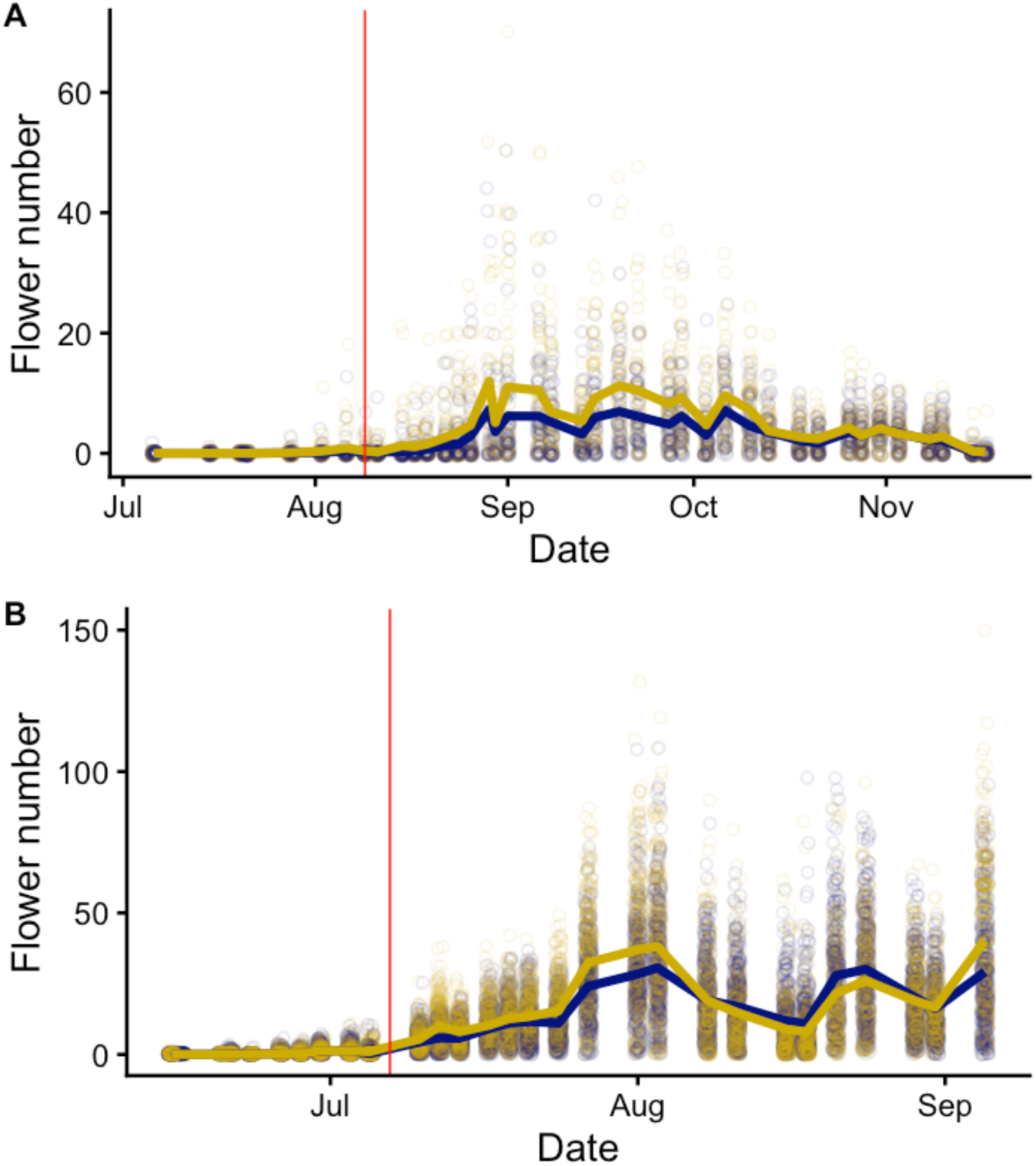
Seasonal flowering dynamics under dicamba drift exposure. Daily flower production for control (blue) and dicamba drift–exposed (yellow) plants in (A) Experiment 1 (2022) and (B) Experiment 2 (2023). Open circles show the number of flowers produced per plant on each census day, and solid lines represent daily mean flower production for each treatment. The red vertical line marks the date on which drift-dose dicamba was applied to treated plants (9 August 2022 in Experiment 1; 7 July 2023 in Experiment 2).

In comparison, in Field Experiment 2 (2023), we found no evidence for a treatment effect on growth (F_1, 676.05_ = 0.02, P = 0.88; Fig. 2E), but like that of Field Experiment 1, we again found that dicamba drift increased the total number of flowers produced across the season (F_1, 336.80_ = 5.35, P = 0.02; Fig. 2F). This effect appeared strongest in the first month post-spray (Fig. 3B), with drift-exposed plants producing ∼29% more flowers, on average, during the first month after exposure (180 vs 140 flowers; F_1, 337.08_ = 22.84, P < 0.001; Fig. 2G, Fig. 3B). We found no evidence that fruit number differed between treatment environments (χ² = 0.64, P = 0.43; Fig. 2H), mirroring our findings examining seed number in the prior experiment.

### Structural equation modeling of reproductive output

We used piecewise structural equation modeling to test whether dicamba exposure influenced reproductive output indirectly through correlated changes in growth and floral allocation across two field experiments (Fig. 4; Table S3). In both years, the best-supported models converged and adequately captured the hypothesized causal relationships based on tests of directed separation and relative model fit.

**Figure 4.**
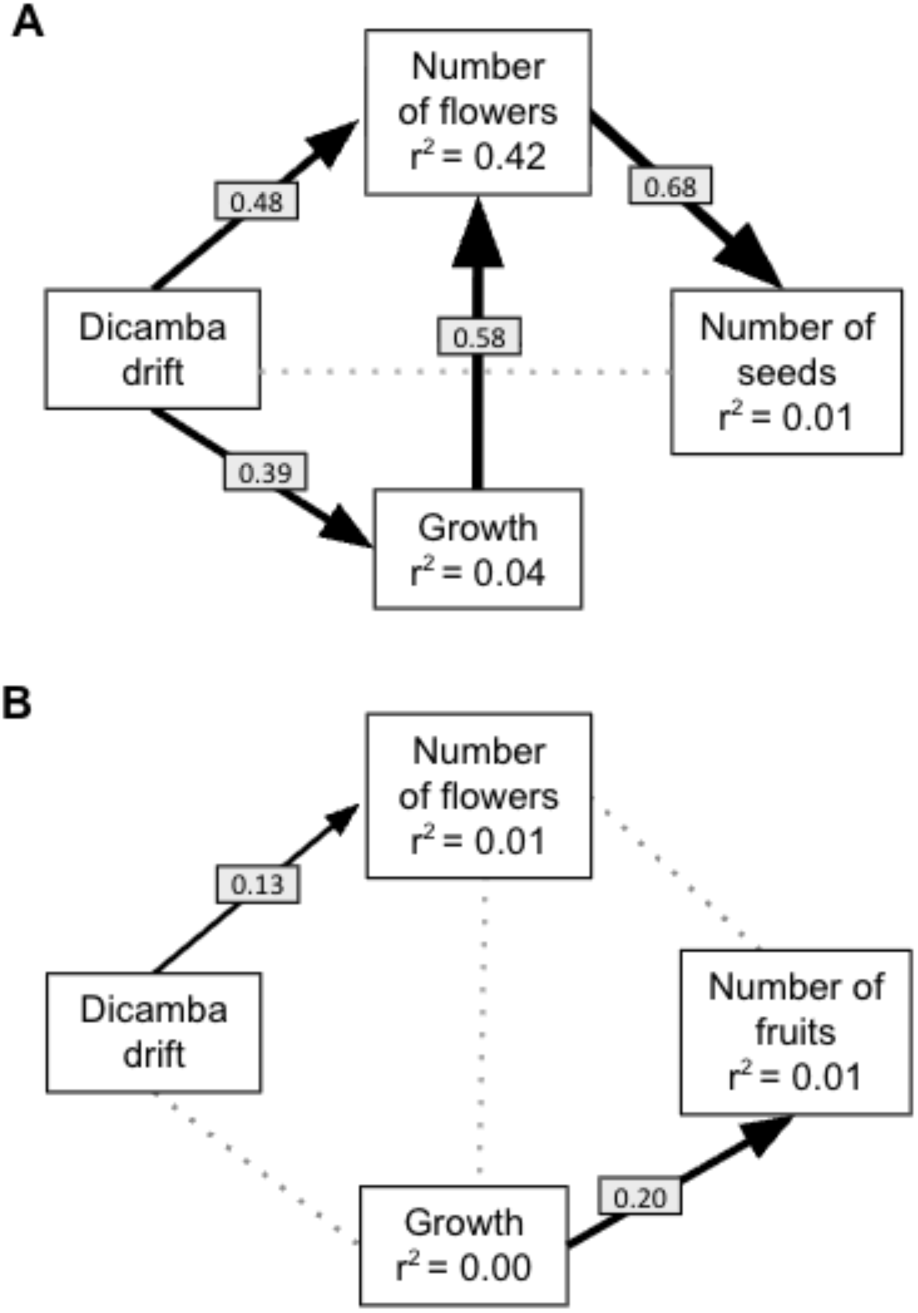
Structural equation models describing the effects of dicamba drift on plant growth and reproduction across two field experiments. Path diagrams show the hypothesized causal relationships among dicamba drift treatment, plant growth, and reproductive output in Field experiment 1 (A) and Field experiment 2 (B). Boxes represent observed variables and arrows indicate directional causal paths. Standardized path coefficients are shown above each arrow; significant paths are displayed as solid black arrows, whereas nonsignificant paths are indicated with grey dotted arrows. The r^2^ values within each response variable denote the proportion of variance explained by the full model. AIC-based comparisons of alternative pSEMs are reported in Table S3. Direct, indirect, and total effects for all pathways are provided in Table S4.

In Experiment 1 (2022; Fig. 4A), dicamba exposure increased reproductive output entirely through indirect effects on floral allocation. Dicamba positively influenced flower number both directly and indirectly *via* enhanced post-spray growth, and flower number was the dominant predictor of seed production. Once flower number was included in the model, neither dicamba nor growth exerted detectable direct effects on seed output. Standardized path coefficients revealed two positive indirect pathways linking dicamba to seed production: a growth-mediated pathway (dicamba -> growth → flowers → seeds; 0.39 × 0.59 × 0.68 = 0.16) and a direct floral allocation pathway (dicamba → flowers → seeds; 0.48 × 0.68 ≈ 0.33). Together, these pathways produced a substantial positive total indirect effect (0.49), indicating that hormetic responses in 2022 were driven by increased floral allocation rather than direct enhancement of seed production.

In Experiment 2 (2023; Fig. 4B), dicamba effects on reproduction were absent. Although dicamba increased flower number, neither flower number nor post-spray growth mediated any effect of dicamba on fruit production. Both indirect pathways linking dicamba to fruit output—*via* flower number (0.22 × 0.02 ≈ 0) and *via* a direct dicamba → fruit pathway (0.01)—were negligible, yielding a total effect of zero. Fruit production was instead associated with growth independently of treatment, and models including dicamba–reproduction pathways were unsupported. Thus, in 2023 dicamba altered floral display without translating into increased reproductive output, highlighting strong context dependence in the expression of hormetic responses.

### Genetic variation in fitness-related traits

We found little evidence for genetic variation in treatment-specific responses in Field Experiment 1 (2022). Neither maternal line nor the maternal line × treatment interaction explained significant variance for growth, flowering phenology, flower number, or seed production (all χ² ≤ 0.19, all P ≥ 0.75 for maternal line; all χ² ≤ 0.28, all P ≥ 0.87 for maternal line × treatment; Table S2). Maternal line variance for seed production was negligible (χ² = 0.10, P = 0.75), providing no evidence for underlying genetic structure in reproductive output in this year.

In contrast, several traits in Field Experiment 2 (2023) showed clear maternal line effects (Table S2). Maternal line significantly explained variation in total flower number (χ² = 14.43, P < 0.001) and early-season flower number (χ² = 10.19, P = 0.001), and showed a marginal effect on fruit production (χ² = 3.45, P = 0.063). These results indicate substantial among-line variation in reproductive traits in 2023. However, the maternal line × treatment interaction was nonsignificant for all traits in both years (all P ≥ 0.23), indicating limited evidence that maternal lines differed in the magnitude of their response to dicamba drift. Collectively, these results provide no evidence for genetic variation underlying hormetic responses in any fitness-related trait examined.

### Floral allocation

Dicamba drift significantly reduced flower width in the 2023 field experiment (Fig. 5A; Table S5). Flowers produced under drift conditions were significantly smaller than those produced in the control environment (F₁,15.8 = 11.41, P = 0.0039), with mean flower width declining from 8.73 mm (95% CI: 8.01–9.47 mm) in control plots to 8.09 mm (95% CI: 7.43–8.77 mm) under dicamba drift, representing an approximate 0.64 mm reduction in size. Importantly, maternal lines differed significantly in their response to dicamba exposure (χ²₂ = 7.46, P = 0.024), providing evidence for genetic variation in tolerance or plasticity of flower width. Population-level variation in flower width was weaker (χ²₁ = 3.05, P = 0.081).

**Figure 5.**
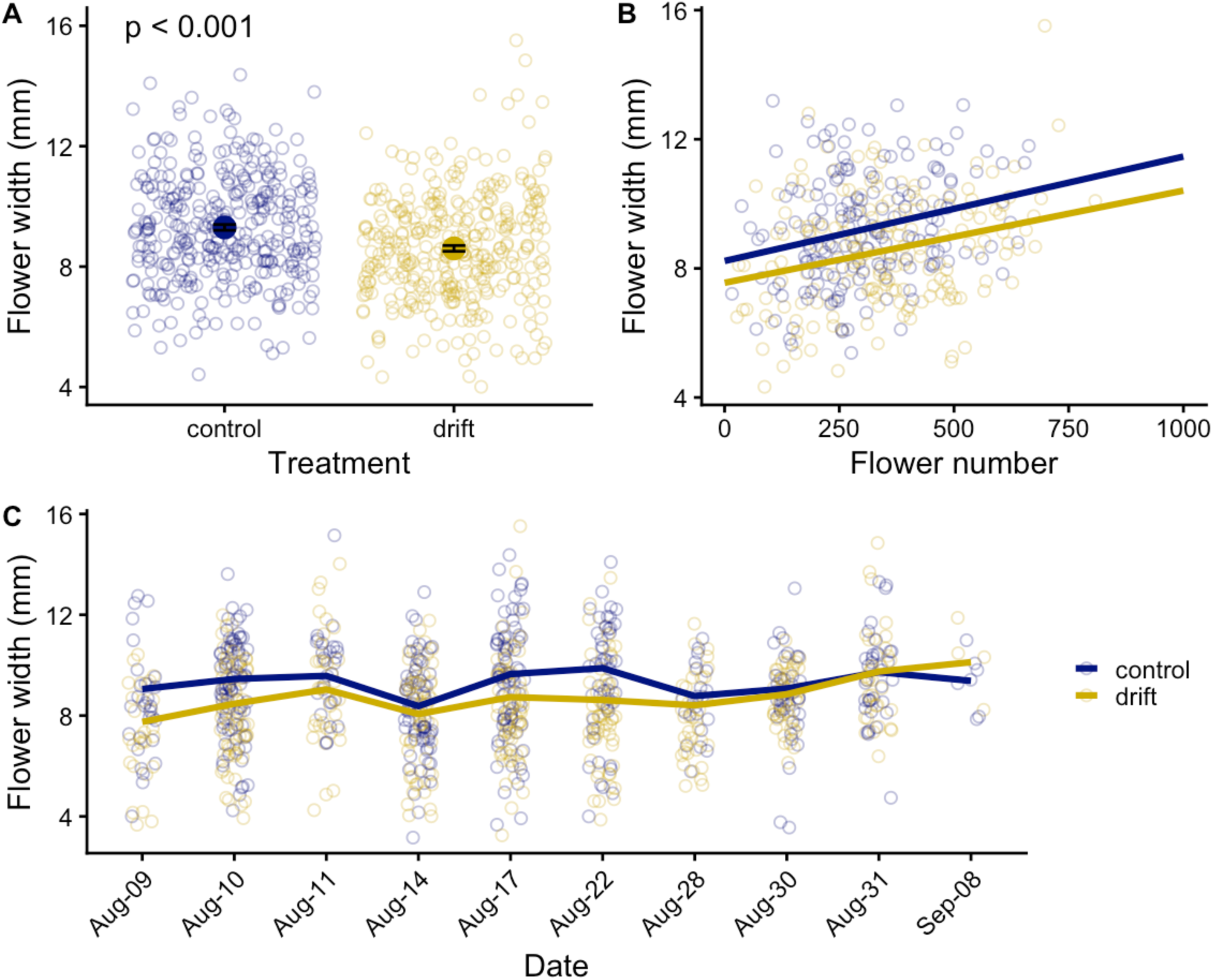
Effects of dicamba drift on flower width and its relationship to reproductive traits. (A) Mean flower width (± SE) for control (blue) and dicamba drift–exposed (yellow) plants, with individual flowers shown as open circles. (B) Relationship between mean flower width and total seasonal flower number, with treatment-specific regression lines. (C) Daily flower width measurements across the flowering period, with individual flowers shown as open circles and smoothed lines depicting temporal trends for each treatment. All data are from Experiment 2.

Despite this reduction in flower size, the slope of the relationship between flower number and flower width did not differ between treatments. Flower width increased significantly with flower number in both environments (χ²₁ = 176.66, P < 0.001; Fig. 5B), and the flower number × treatment interaction was not significant (χ²₁ = 1.19, P = 0.28), indicating that dicamba drift did not alter how floral display size scaled with flower abundance. Thus, although individual flowers were smaller in the drift environment, plants did not shift their allocation strategy according to flower number.

The reduction in flower size persisted across sampling dates. When daily flower width measurements were plotted over the month following dicamba application (Fig. 5C), drift-treated plants produced smaller flowers throughout the sampling period, with no evidence of convergence toward control values. Flower size changes were parallel between treatments, further reinforcing that dicamba drift consistently reduced per-flower investment over time.

### Pollinator observation

Pollinator visitation rate differed significantly by treatment (χ²₁ = 9.79, P = 0.0018; Fig. 6A, Table S6), with lower visitation under dicamba drift (0.0278 visits per flower per minute; 95% CI: 0.020–0.039) than in control plants (0.0415; 95% CI: 0.030–0.057). This corresponds to an approximately 33% reduction in visitation under drift exposure, with pollinators visiting control flowers 1.49 times more frequently than drift-exposed flowers (control/drift ratio = 1.49).

**Figure 6.**
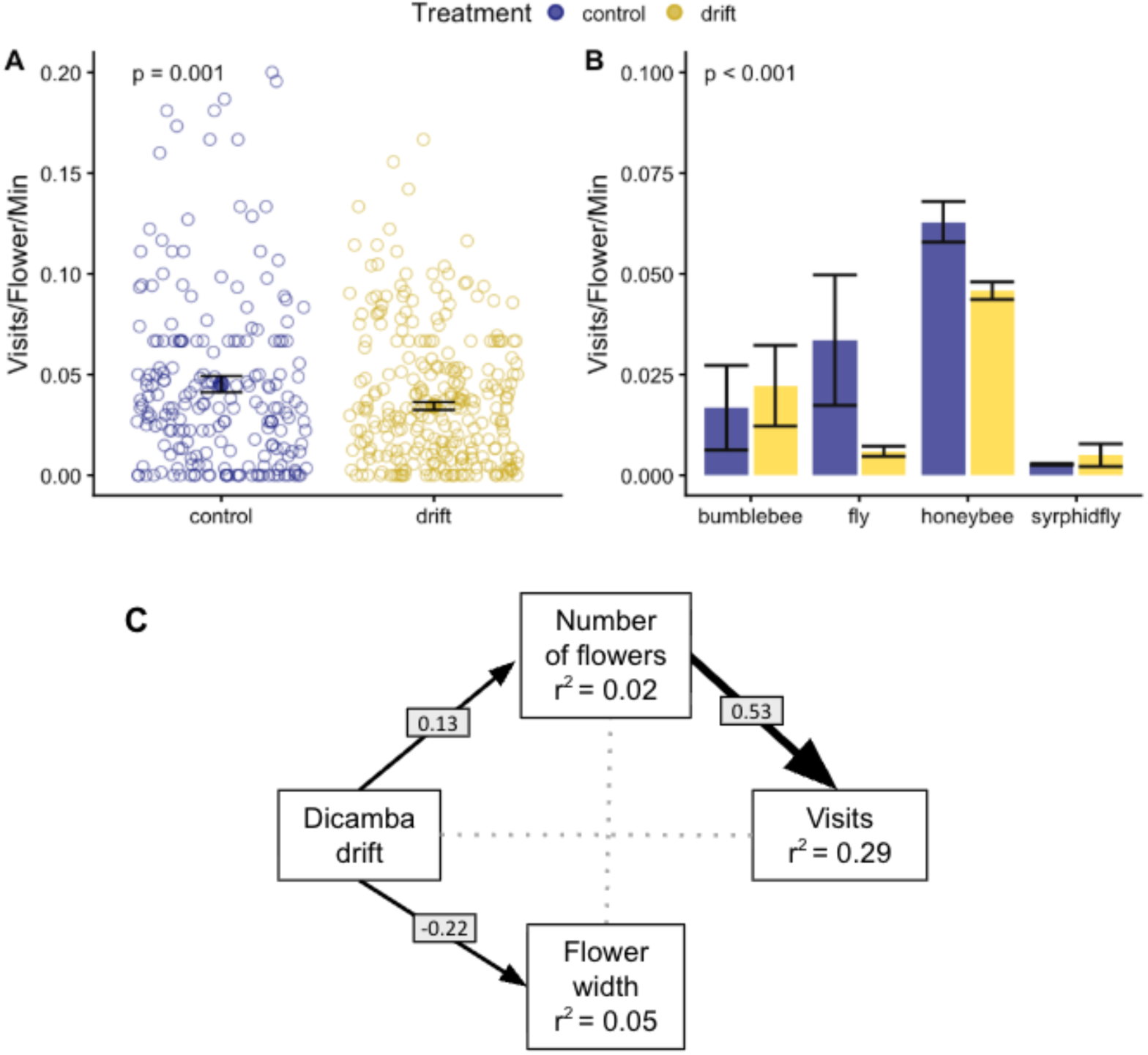
Dicamba drift alters pollinator visitation indirectly through changes in flower number. (A) Pollinator visitation rate (visits per flower per minute) under control and dicamba drift treatments. Points show individual observations; horizontal bars indicate treatment means ± SE. (B) Mean visitation rates by pollinator functional group under control and dicamba drift treatments (± SE). P-value reflects the functional group x treatment interaction. (C) Structural equation model summarizing direct and indirect effects of dicamba drift on pollinator visitation. Standardized pathway coefficients are shown next to arrows; arrow thickness is proportional to effect size. The direct effect of dicamba drift on visitation (dotted arrow) and pathways involving flower width were not significant. The r^2^ values within each response variable denote the proportion of variance explained by the full model.

Pollinator visitation differed among functional groups and responded to dicamba exposure in a group-specific manner (Treatment × functional group interaction: χ²₃ = 18.77, P < 0.001; Fig. 6B). Visitation by flies and honeybees was significantly reduced under dicamba drift, with flies visiting control flowers more than four times as frequently as drift-exposed flowers (control/drift ratio = 4.32, P < 0.001) and honeybee visitation reduced by approximately 33% under drift exposure (control/drift ratio = 1.49, P = 0.002). In contrast, bumblebee visitation increased slightly under drift (P = 0.049), while syrphid fly visitation did not differ between treatments (P = 0.46).

### Structural equation modeling of pollinator visitation

We performed structural equation modeling to determine which component of floral display—flower number or flower size—most strongly influenced pollinator visits, quantified as visits per minute. We compared alternative structural equation models to evaluate whether dicamba effects on pollinator visits were mediated primarily through changes in floral display size or individual flower size, using a fully saturated model as a reference. The best-supported model, which fit the data well (Fisher’s C = 0.13, df = 2, P = 0.94), indicated that dicamba influenced pollinator visitation indirectly through effects on floral traits rather than through a direct pathway. Specifically, dicamba significantly reduced flower width (β = −0.22, P < 0.001) while increasing flower number (β = 0.13, P = 0.021), and flower number had a strong positive effect on pollinator visitation time (β = 0.53, P < 0.001). In contrast, dicamba had no detectable direct effect on visitation after accounting for floral traits (β = −0.10 ± 0.10 SE, P = 0.31), and flower width did not significantly predict either flower number (P = 0.37) or pollinator visitation; accordingly, models including a direct flower width–to–visitation pathway were less well supported (ΔAIC ≈ 2). The resulting indirect effect of dicamba on visitation mediated by flower number was positive (β indirect ≈ 0.07). This apparent contrast reflects differences in scale between visitation metrics, with SEM capturing whole-plant visitation rather than per-flower rates.

## Discussion

In this study, we examined herbicide-induced hormesis through the lens of tolerance and overcompensation using the weed *Oxalis stricta* as a model system. Across two field experiments, exposure to drift-relevant doses of dicamba elicited changes in growth, flowering, and allocation traits, but the magnitude, direction, and downstream consequences of these responses varied between years. By modeling causal relationships among growth, reproductive traits, and biotic interactions using structural equation modeling, we show that hormetic responses primarily reshape trait expression and allocation patterns, with fitness consequences that are context dependent. In some cases, these trait shifts propagated indirectly to reproductive output, whereas in others they altered floral display and plant–pollinator interactions without translating into increased fitness. These findings place herbicide-induced hormesis within an evolutionary ecology framework, highlighting how low-dose chemical stress can reshape trait expression and fitness, even when limited G×E variation suggests little potential for evolutionary change.

### Hormesis as compensatory flowering under herbicide drift

Our prior work showed that *Oxalis stricta* does not survive or maintain seed production when exposed to the recommended field dose of dicamba (Baucom *et al*., 2025), indicating that this species is neither resistant nor tolerant to this herbicide at agronomic rates. In contrast, flowering responses under drift-level dicamba exposure were consistently positive. Across both prior studies and the present experiments, dicamba drift increased flower number and, in some cases, altered flowering timing or duration relative to non-exposed controls (Iriart *et al*., 2022; Baucom *et al*., 2025). Extending these observations, our two field experiments provide clear evidence of compensatory responses: in both years, dicamba exposure consistently increased flowering, whereas effects on downstream components of fitness were more variable. In 2022, enhanced flowering was accompanied by increased vegetative growth and propagated indirectly to higher seed production, whereas in 2023 flowering increases did not translate into greater fruit output. This contrast indicates that flowering is a robust hormetic response to dicamba drift in *O. stricta*, while the fitness consequences of that response depend strongly on environmental or developmental context.

One likely contributor to these year-specific outcomes is the timing of herbicide exposure relative to plant development. In the first experiment, dicamba was applied after 22 days of growth, when only 17% of plants had initiated flowering, whereas in the second experiment 78% of individuals had flowered within that same interval, such that plants were at a more advanced reproductive stage at the time of exposure. Developmental timing has been shown to shape the expression of hormesis in other plant systems (Velini *et al*., 2008; De Carvalho *et al*., 2013; Belz & Duke, 2014), with earlier exposure often eliciting stronger compensatory growth or reproductive responses. Although both experiments were conducted in the same field location, additional year-to-year variation in environmental conditions (*e.g*., temperature, precipitation) may also have influenced which traits most strongly determined fitness in a given year. Together, these factors likely explain why consistent increases in flowering were associated with higher seed production through indirect trait pathways in 2022, but not in 2023, when increased flowering did not translate into detectable differences in reproductive output. Such variability is common in hormesis studies and has complicated efforts to predict or exploit low-dose herbicide responses in agricultural systems (Duke *et al*., 2025). Future experiments that explicitly manipulate exposure timing and environmental context will be essential for resolving how and when compensatory flowering responses lead to fitness gains.

Further, although we have not yet conducted a full dose-response experiment, the contrast between field-dose lethality and drift-dose survival, combined with increased flowering across years, offers compelling evidence that hormesis operates in this species and underscores the importance of evaluating herbicide drift effects under ecological field conditions. Our findings align with the broader literature: while dicamba-specific hormesis studies in weedy plants remain limited, our results parallel hormetic responses to sublethal doses of herbicides such as glyphosate and ALS inhibitors (summarized in (Belz *et al*., 2022; Duke *et al*., 2025).

Our use of structural equation modeling to examine hormesis is, to our knowledge, novel, providing a framework to evaluate how low-dose herbicide exposure influences fitness through interconnected traits. By explicitly modeling causal relationships among growth, flowering, and reproductive output, we were able to distinguish consistent trait responses from year-specific fitness outcomes. In 2022, dicamba had no detectable direct effect on seed production; instead, fitness gains emerged indirectly through a cascade of trait responses in which enhanced vegetative growth increased flower production, which in turn drove higher seed output. In contrast, although dicamba consistently increased flowering in 2023, neither flower number nor growth translated into increased fruit production, and no direct treatment effect on reproduction was supported. These contrasting pathway structures indicate that while flowering responses to dicamba drift are robust, their consequences for fitness depend on which traits most strongly limit reproduction in a given year.

Taken together, the SEM results demonstrate that dicamba exposure reshaped growth and allocation traits in both experiments, but only under certain ecological or developmental contexts did these changes propagate to reproductive output. This strong interannual context dependence underscores the value of multivariate, causal approaches for understanding hormesis, revealing how low-dose chemical stress can reorganize trait relationships even when its effects on fitness are inconsistent across environments.

### No evidence for genetic variation in hormetic responses

Despite consistent effects of dicamba on flowering and other allocation traits, we found little evidence for genetic variation underlying hormetic responses in any fitness-related trait. Maternal line × treatment interactions were nonsignificant in both years, indicating that lines did not differ in their plastic responses to dicamba drift and suggesting limited potential for the evolution of increased hormesis in *Oxalis stricta*. Similar patterns have been reported in other systems: *Trifolium pratense* exposed to synthetic auxin drift likewise detected strong trait-level responses but little evidence for line × treatment interactions in growth-related traits, consistent with predominantly plastic responses to sublethal exposure (Iriart *et al*., 2025). In contrast, studies in barley (*Hordeum vulgare*; (Belz & Sinkkonen, 2019, 2021) and weedy plants *Tripleurospermum perforatum* (Belz & Sinkkonen, 2016) and *Abutilon theophrasti* (Ethridge *et al*., 2023) strongly suggest that genetic variation in hormetic responses can occur at low herbicide doses.

In contrast to the absence of genetic variation in hormesis, we did detect significant among-line variation in reproductive traits such as flower number, indicating ample genetic variation in baseline fitness traits themselves. Thus, while reproductive performance may evolve, the plastic responses to dicamba drift exposure likely will not, and hormesis is unlikely to drive adaptive change in this species. While we detected little evidence for genetic variation in responses at the drift dose used, we cannot exclude the possibility that genetic variation in hormesis may emerge at other sublethal exposure levels or under different environmental conditions. Thus, an important next step will be to evaluate hormetic responses across multiple sublethal doses, as recommended by (Duke *et al*., 2005), to determine whether genetic variation emerges at particular dose thresholds, which would alter expectations about the evolutionary trajectory of hormesis in this species.

### Dicamba drift reshapes floral allocation and display traits

Given the absence of genetic variation underlying hormetic responses in *Oxalis stricta*, these responses are unlikely to evolve, and their consequences are more likely to be expressed through altered ecological interactions rather than adaptive change in *Oxalis*. Across both field experiments, dicamba drift consistently increased floral display, reinforcing prior evidence that sublethal exposure promotes elevated flower production in this species (Iriart *et al*., 2022). We therefore asked whether this increase in flower number was accompanied by shifts in floral allocation. Consistent with this expectation, dicamba exposure significantly reduced flower width throughout the flowering period, even as flower production increased. This pattern suggests a compensatory allocation strategy in which plants produce more flowers at the expense of per-flower investment, resulting in a larger but finer-scaled floral display. Such size reductions have important ecological implications, as floral morphology strongly influences pollinator attraction and pollen transfer efficiency (Caruso *et al*., 2019); persistent declines in flower size under drift could alter pollination dynamics and, potentially, patterns of mating and gene flow. Similar patterns of altered floral display under dicamba drift have been documented in other species, including shifts in flower number, phenology, and pollinator visitation (Egan *et al*., 2014; Bohnenblust *et al*., 2016; Baucom *et al*., 2025), indicating that auxin-mimicking herbicides can restructure floral traits in ways that extend beyond simple biomass responses.

### Consequences for plant–pollinator interactions

Consistent with these allocation shifts, dicamba drift significantly altered pollinator visitation patterns, indicating that low-dose herbicide exposure reshapes plant–pollinator interactions rather than uniformly suppressing pollinator activity. Overall visitation rates declined under drift exposure, but responses differed strongly among functional groups, revealing a treatment × pollinator interaction rather than a generalized reduction in visitation. Visitation by flies and honeybees declined substantially in the drift environment, whereas bumblebee visitation increased slightly and syrphid fly visitation was unaffected. These group-specific responses suggest that pollinators respond differently to dicamba-induced changes in floral display, likely reflecting variation in foraging strategies and sensitivity to display size rather than uniform avoidance of drift-exposed plants.

Structural equation modeling clarified the potential mechanistic basis of these patterns. Dicamba had no detectable direct effect on pollinator visitation once floral traits were included, nor did changes in flower width contribute to variation in visitation. Instead, dicamba influenced visitation predominantly through its hormetic effect on flower number, which had a modest but positive indirect effect on visit rate. This apparent contrast with the mixed-effects results reflects differences in scale and response metrics: mixed-effects models quantified visitation as visits per flower per minute, whereas the SEM evaluated visitation at the plant level, integrating the expanded floral display (*i.e.,* visits/minute rather than visits/flower/minute). Together, these results indicate that dicamba-exposed plants receive similar or slightly greater total pollinator attention due to increased flower production (as summarized by Fig 6C), but that this attention is distributed across more flowers, resulting in reduced visitation rate per flower (evident in Fig 6A). Thus, dicamba drift alters pollinator behavior primarily by redistributing pollinator effort across an increased floral display rather than through changes in individual flower size, with potentially important consequences for pollen transfer efficiency and mating outcomes.

### Implications for weed ecology and eco-evolutionary dynamics

Overall, our findings align with the tolerance–overcompensation framework (Fig. 1) but reveal important context dependence in how hormetic responses translate to fitness. Across both experiments, *Oxalis stricta* consistently increased flower production under dicamba drift, indicating a compensatory allocation response rather than a uniform increase in reproductive success. However, this enhanced floral allocation resulted in overcompensation—defined as increased fitness under stress—only in 2022, when elevated flower production indirectly led to higher seed output. In contrast, in 2023 increased flowering did not translate into increased reproduction, underscoring that compensatory trait responses do not necessarily imply overcompensation in fitness. Because we detected no heritable variation in the magnitude of these responses, hormesis in *O. stricta* appears to reflect plastic tolerance rather than an evolving overcompensatory strategy.

While our study focuses on a single species and herbicide, the trait-mediated pathways identified here are likely relevant to other non-target plants exposed to synthetic auxin drift. The patterns we document are especially timely given recent shifts in U.S. agriculture, where the widespread adoption of auxin-resistant crop varieties has driven substantial increases in the use of synthetic auxin herbicides such as dicamba and 2,4-D since the early 2000s (Singh & Jhala, 2025). Although regulatory changes have recently redirected use away from dicamba (Singh & Jhala, 2025), auxin drift remains a widespread and emerging feature of agricultural landscapes, particularly with the predicted increase in 2,4-D use in the coming decade (Freisthler *et al*., 2022).

This shift toward synthetic auxin herbicides is ecologically consequential because auxins are central regulators of plant development (Woodward & Bartel, 2005) and influence interactions with herbivores and pollinators through effects on defense chemistry, volatile signaling, and floral traits (Ramos *et al*., 2021, 2023; Johnson *et al*., 2023). Accordingly, synthetic auxin drift has the capacity to reshape not only plant growth and reproduction but also plant–insect interactions. The combination of increased floral display and reduced flower size under drift that we document in *O. stricta* suggests that low-dose exposure may alter traits important for pollinator attraction, pollen transfer efficiency, and mating outcomes, even in the absence of consistent fitness overcompensation.

These responses have implications for weed dynamics in agroecosystems. Plastic, compensatory increases in reproduction may promote persistence under low-dose exposure, favoring species capable of such responses and potentially reshaping weed community composition. Likewise, auxin-induced modification of floral traits may influence pollination networks and gene flow even in the absence of evolutionary change in hormesis itself (Iriart *et al*., 2022). Future work should evaluate whether these plastic responses affect pollinator visitation, seedling recruitment, and longer-term population trajectories, and whether drift exposures of synthetic auxins do in fact facilitate weed shifts.

## Acknowledgements

We thank Mike Palmer and Jeremy Moghtader and Matthaei Botanical Gardens for assistance with field preparation and maintenance. We thank Betania M Cornejo, Yasmina Zimmer, Julie Sheldon, and Robin Pitilon for help with data collection and entry and thank Grace Zhang and the Baucom lab for feedback on the manuscript. This work was supported by the University of Michigan.

## Author contributions

AS and RSB designed the research; AS, EK, and OT collected data; AS and RSB analyzed the data; AS and RSB interpreted results and wrote the manuscript.

## Competing interests

The authors declare no competing interests.

## Data availability

Data will be made available on Dryad upon acceptance.

---

**Table S1:**
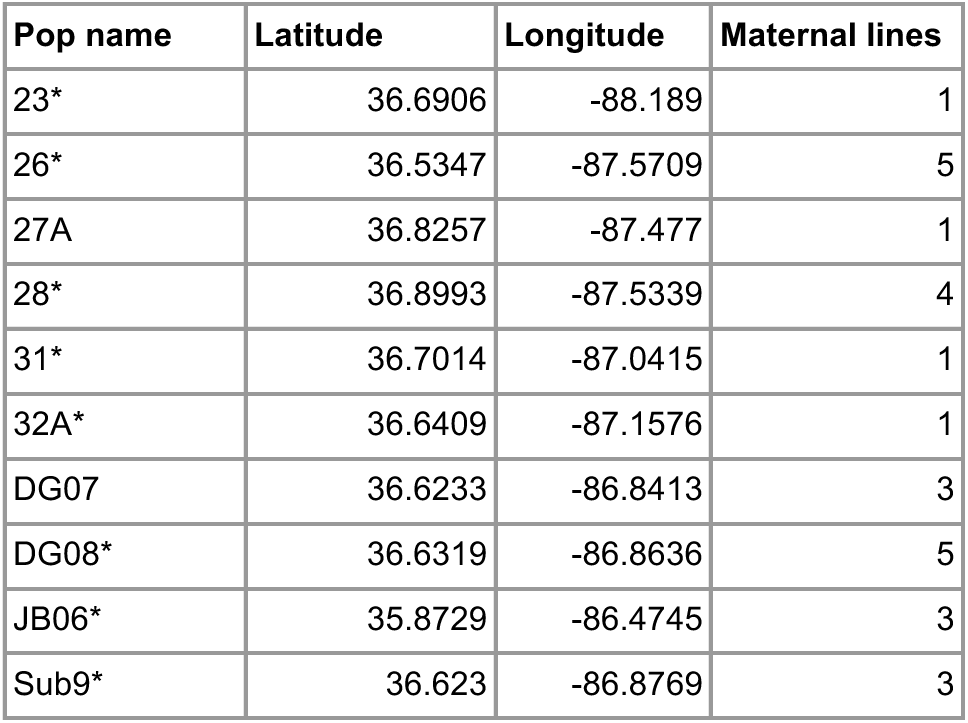
Locations of the 10 populations from which seeds were sampled. Starred populations are represented again in the 2023 experiment. Maternal lines are defined as individual plants from which seeds were collected in the field and then selfed to plant for the experiment.

**Figure S1.**
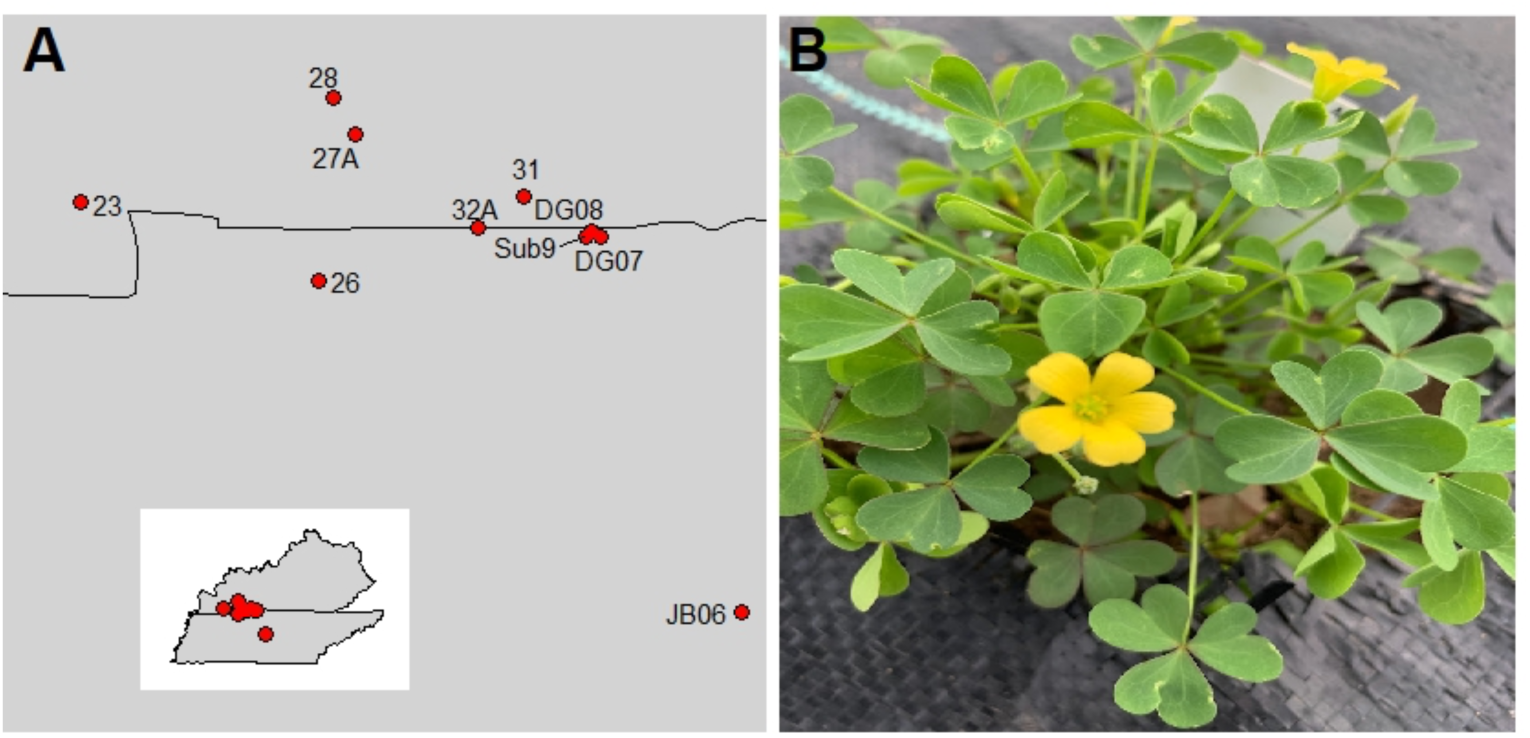
*Oxalis stricta* sampling locations and species image. A) A map of the collection locations for each population in the field experiment. Locations were chosen based on proximity to farm field and presence of the species. B) A photo of *Oxalis stricta* in the field. Over the course of a season, it will produce many yellow primarily selfing flowers and dehiscent exploding fruits.

**Table S2.** Mixed-effects model results for Experiments 1 (2022) and 2 (2023). The table reports fixed-effect tests for dicamba treatment across vegetative growth, flowering, and reproductive traits in Experiment 1 (2022) and Experiment 2 (2023), along with likelihood ratio tests for associated random effects. Prespray size covariates were included where relevant.

| **A. Experiment 1 - 2022** |  |  | **Growth** |  |  | **Date of first flower** |  |  | **Number flowers** |  |  | **Number seeds** |  |  |
| --- | --- | --- | --- | --- | --- | --- | --- | --- | --- | --- | --- | --- | --- | --- |
| *Fixed effects* |  |  | df | F | P | df | F | P | df | F | P | df | chisq | P |
| Treatment |  |  | 1, 171.13 | 4.616 | **0.033** | 1, 106 | 0.504 | 0.479 | 1, 121.01 | 12.229 | **0.001** | 1 | 0.58 | 0.446 |
| Pre-spray leaf number |  |  | -- | -- | -- | 1, 106 | 70.588 | **<0.001** | 1, 127.22 | 37.771 | **<0.001** | 1 | 5.039 | **0.025** |
| *Random effects* |  |  | df | chisq | P | df | chisq | P | df | chisq | P | df | chisq | P |
| Maternal line |  |  | 1 | 0 | 1 | 1 | 0 | 1 | 1 | 0.102 | 0.75 | 1 | 0.104 | 0.748 |
| Maternal line x treatment |  |  | 3 | 0 | 1 | 2 | 0 | 1 | 1 | 0.186 | 0.911 | 2 | 0.283 | 0.868 |
| Population |  |  | 1 | 1.832 | 0.176 | 1 | 0 | 1 | 1 | 1.049 | 0.306 | 1 | 0 | 1 |
| Block |  |  | 1 | 0.225 | 0.636 | -- | -- | -- | -- | -- | -- | -- | -- | -- |
| **B. Experiment 2 - 2023** |  |  | **Growth** |  |  | **Number flowers, overall** |  |  | **Number flowers, 1st month post-spray** |  |  | **Number fruits** |  |  |
| *Fixed effects* |  |  | df | F | P | df | F | P | df | F | P | df | chisq | P |
| Treatment |  |  | 1, 676.05 | 0.021 | 0.883 | 1, 336.80 | 5.349 | **0.021** | 1, 337.08 | 22.839 | **<0.001** | 1 | 0.635 | 0.426 |
| Pre-spray leaf number |  |  | -- | -- | -- | 1, 339.02 | 0.723 | **<0.001** | 1, 334.98 | 2.764 | 0.0974 | 1 | 24.922 | **<0.001** |
| *Random effects* |  |  | df | chisq | P | df | chisq | P | df | chisq | P | df | chisq | P |
| Maternal line |  |  | 1 | 0 | 1 | 1 | 14.432 | **<0.001** | 1 | 10.185 | **0.001** | 1 | 3.449 | 0.063 |
| Maternal line x treatment |  |  | 3 | 0 | 1 | 3 | 1.842 | 0.6058 | 3 | 2.512 | 0.473 | 2 | 2.977 | 0.226 |
| Population |  |  | 1 | 1.547 | 0.214 | 1 | 0 | 1 | 1 | 0 | 1 | 1 | 4.349 | 0.037 |
| Block |  |  | 1 | 0.2502 | 0.617 | 1 | 0 | 0.9999 | 1 | 0 | 1 | 1 | 8.089 | 0.004 |

**Supplemental methods 1**

**Structural equation model selection—reproductive output.**

To evaluate whether dicamba exposure influenced reproductive output directly or indirectly through correlated growth and allocation traits, we used piecewise structural equation modeling (SEM). For each year, we compared a small set of biologically motivated SEMs that differed in whether reproductive output was assumed to be mediated primarily by flower number, post-spray growth, or both, and whether direct effects of dicamba on reproduction were included. Model adequacy was evaluated using (i) tests of directed separation to identify missing paths, (ii) summed AIC values across component models to compare relative support, and (iii) the significance and magnitude of individual path coefficients. Standardized effects for both seed and flower number were calculated manually on the link scale.

In 2022, the best-supported model included an indirect pathway from dicamba exposure to seed number mediated through increased flower number, with no remaining direct effect of dicamba once flower number was included. In contrast, in 2023, models in which fruit production was mediated primarily by post-spray growth were consistently favored, whereas flower number had little explanatory power for fruit output. We also examined the AIC value of the full, saturated SEMs for comparison, with the general structure of saturated models shown below.

**Table S3.**
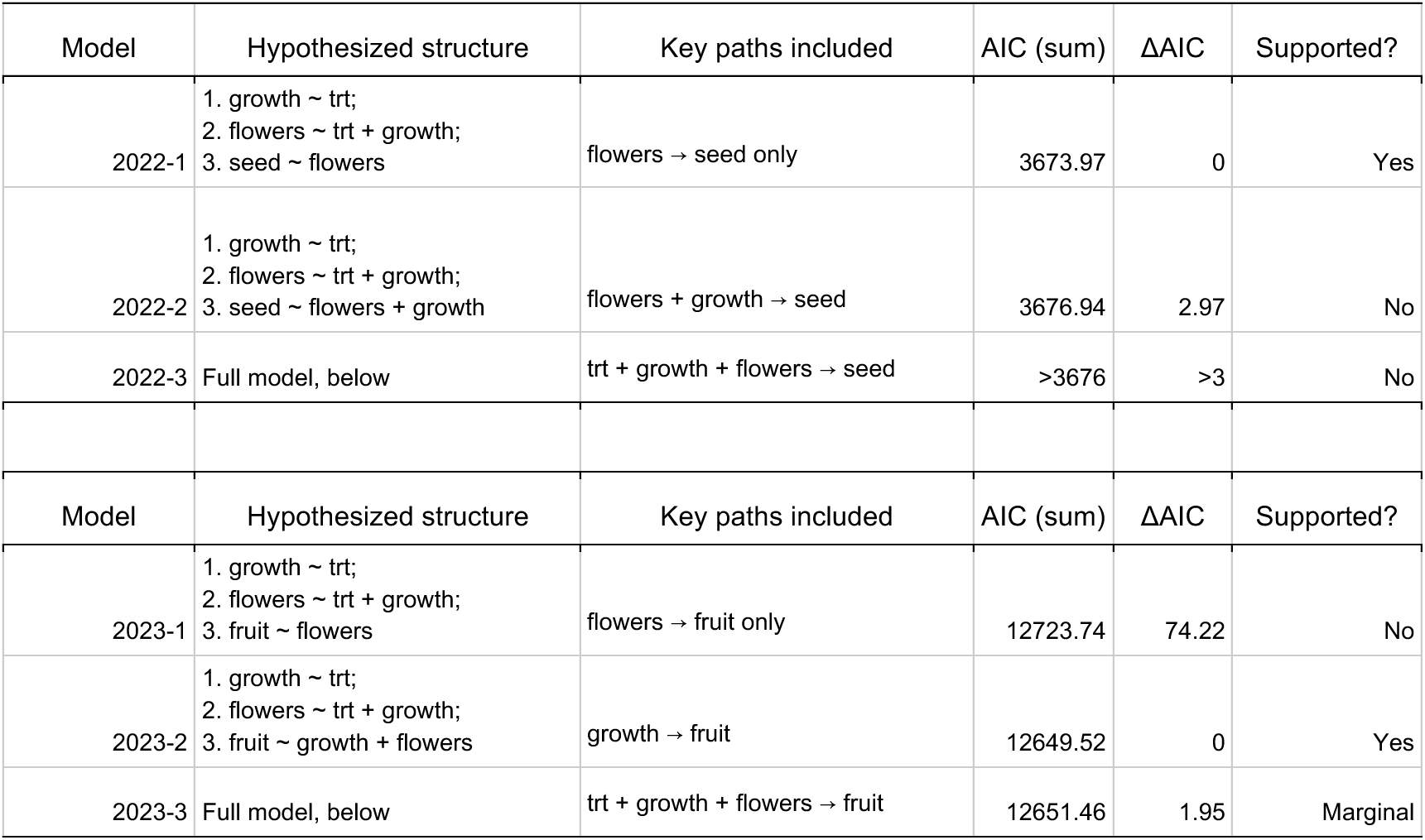
Comparison of piecewise structural equation models (pSEMs) evaluating alternative pathways linking dicamba treatment (trt), post-spray growth, floral allocation, and reproductive output in Field Experiments 1 (2022) and 2 (2023). For each model, we report the hypothesized structure, summed AIC across component models, ΔAIC relative to the best-supported model, and model support.

**Figure S2.**
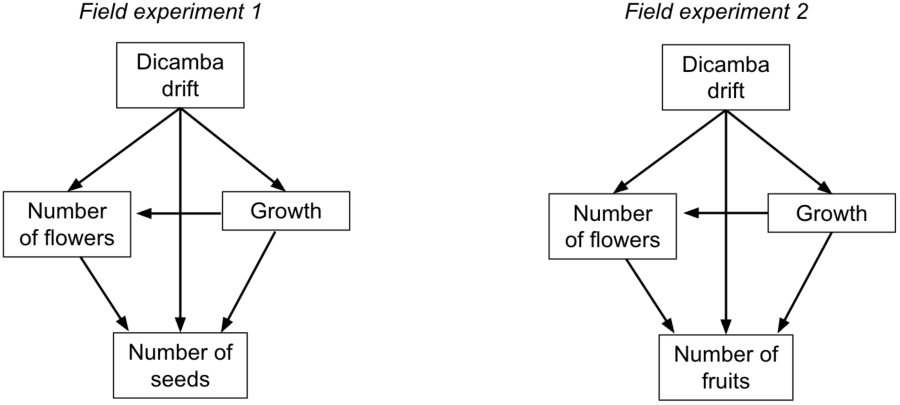
Conceptual path diagrams illustrating the full saturated structural equation models evaluated for comparison with reduced models in Field experiment 1 (2022) and Field experiment 2 (2023).Diagrams show hypothesized direct effects of dicamba drift on growth, reproductive traits, and final fitness components (seed or fruit number), as well as interrelationships among growth and flowering variables included in the saturated models.

**Table S4.** Indirect, direct, and total effects of dicamba drift on fitness from the structural equation models in Fig. 4. Standardized indirect and direct effects of dicamba drift on reproductive output for (A) Experiment 1 (2022) and (B) Experiment 2 (2023), calculated from the pathways in Fig. 4. Indirect effects were obtained by multiplying the standardized coefficients along each causal path; total effects are the sum of all pathways for each experiment.

| A. Experiment 1 |  |  |
| --- | --- | --- |
| Pathway | Description |  |
| 1 | Trt --> Growth --> Num. flowers --> Num. seeds;<br>0.39 x 0.59 x 0.68 | 0.16 |
| 2 | Trt --> Num. flowers --> Num. seeds;<br>0.48 x 0.68 | 0.33 |
|  | Total effect | 0.49 |
| B. Experiment 2 |  |  |
| Pathway | Description |  |
| 1 | Trt --> Num. flowers --> Num. fruits;<br>0.22 x 0.02 | 0 |
| 2 | Trt --> Num. fruits; 0.01 | 0 |
|  | Total effect | 0 |

Effects are calculated as products of standardized path coefficients from the best-supported piecewise structural equation models for each experiment. For 2022, the reduced SEM was supported, and treatment effects on seed production were entirely indirect via growth and flower number. For 2023, the best-supported SEM included growth → fruit but no treatment → fruit or flower → fruit paths; values of 0.00 reflect negligible or unsupported pathways rather than true zeros.

**Table S5.** Results of mixed-effects models testing the effects of dicamba treatment on average flower width. Fixed effects were evaluated using F tests, and random effects were assessed using likelihood ratio tests (χ²). Degrees of freedom, test statistics, and P values are reported.

|  | Flower width |  |  |
| --- | --- | --- | --- |
| *Fixed effects* | df | F | P |
| Treatment | 1, 15.81 | 11.41 | 0.003 |
| *Random effects* | df | chisq | P |
| Maternal line | 1 | 12.99 | <0.001 |
| Maternal line x treatment | 1 | 7.46 | 0.024 |
| Population | 1 | 3.05 | 0.081 |

**Table S6.**
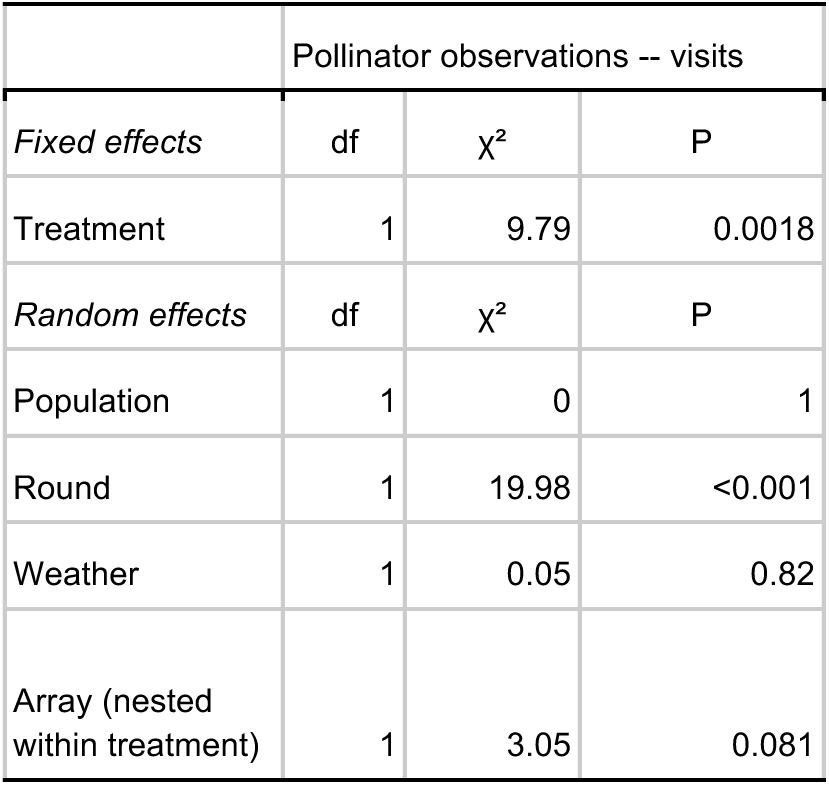
Results of mixed-effects models testing the effects of dicamba treatment on pollinator visitation rate (visits per flower per minute). Fixed effects were evaluated using Type III Wald χ² tests, and random effects were assessed using likelihood ratio tests; array was modeled as a random effect nested within treatment.

**Supplemental Methods 2**

**Structural equation model selection**—To evaluate whether dicamba exposure influenced pollinator visitation directly or indirectly through changes in floral traits, we used piecewise structural equation modeling (SEM). We compared a small set of biologically motivated SEMs that differed in how flower width and flower number were assumed to influence one another and pollinator visitation, and in whether direct effects of dicamba on visitation were included. In all models, dicamba treatment was specified as an exogenous driver with potential direct effects on flower width, flower number, and pollinator visit time. Models were evaluated using (i) tests of directed separation to identify missing paths, (ii) summed AIC values across component models to compare relative support, and (iii) the significance and magnitude of individual path coefficients. For the non-Gaussian visitation component, predictor-standardized effects were calculated on the link scale to facilitate comparison of effect sizes.

We evaluated three piecewise structural equation models representing distinct hypotheses about how dicamba-induced changes in floral traits influence pollinator visitation. The first model treated flower width and flower number as parallel traits, modeled as independent responses to dicamba exposure, with pollinator visitation mediated by flower number alone. This model tested whether dicamba effects on visitation were expressed primarily through changes in floral display size, independent of individual flower size. The second model introduced a directional pathway from flower width to flower number, testing whether changes in individual flower size structured floral display size and thereby influenced visitation. The third model was the fully saturated SEM, which additionally allowed flower width to exert a direct effect on pollinator visitation and included all possible paths among variables; this model was evaluated as a reference rather than as a competing candidate structure.

**Table S7.** Comparison of alternative piecewise structural equation models (pSEMs) evaluating hypothesized pathways linking dicamba treatment, flower width, flower number, and pollinator visitation time. For each candidate model, we report the summed AIC across component models, ΔAIC relative to the best-supported model, and model support, with ΔAIC = 0 indicating the best-supported structure, ΔAIC ≤ 2 considered marginal, and ΔAIC > 2 considered unsupported.

| Model | Hypothesized structure | Key paths included | AIC (sum) | $\Delta AIC$ | Supported? |
| --- | --- | --- | --- | --- | --- |
| Poll-1 | 1. flower width ~ trt;<br>2. flowers ~ trt;<br>3. visit time ~ trt + flowers | flowers → visit time only<br>(parallel traits; no width → flowers) | 1995.13 | 0 | Yes |
| Poll-2 | 1. flower width ~ trt;<br>2. flowers ~ trt + width;<br>3. visit time ~ trt + flowers | width → flowers → visit time | 1996.33 | 1.2 | Marginal |
| Poll-3 | Full, saturated model | trt + width + flowers → visit time | 1998.33 | 3.2 | No |

**Table S8.** Decomposition of total treatment effects on pollinator visitation time based on the best-supported piecewise structural equation model. Individual indirect pathways are shown with their component standardized coefficients.

| Pathway | Description | Effect |
| --- | --- | --- |
| 1 | Trt → Num. flowers → Visit time; $0.13 \times 0.5303$ | 0.07 |
|  | Total effect | 0.07 |

## Notes

### Competing Interest Statement

The authors have declared no competing interest.

